# Mammalian cells-based platforms for the generation of SARS-CoV-2 virus-like particles

**DOI:** 10.1101/2023.07.24.550415

**Authors:** Ghada Elfayres, Ricky Raj Paswan, Laura Sika, Marie-Pierre Girard, Soumia Khalfi, Claire Letanneur, Kéziah Milette, Amita Singh, Gary Kobinger, Lionel Berthoux

## Abstract

Severe acute respiratory syndrome coronavirus 2 (SARS-CoV-2) is the causative agent of COVID-19. Though many COVID-19 vaccines have been developed, most of them are delivered via intramuscular injection and thus confer relatively weak mucosal immunity against the natural infection. Virus-Like Particles (VLPs) are self-assembled nanostructures composed of key viral structural proteins, that mimic the wild-type virus structure but are non-infectious and non-replicating due to the lack of viral genetic material. In this study, we efficiently generated SARS-CoV-2 VLPs by co-expressing the four SARS-CoV-2 structural proteins, specifically the membrane (M), small envelope (E), spike (S) and nucleocapsid (N) proteins. We show that these proteins are essential and sufficient for the efficient formation and release of SARS-CoV-2 VLPs. Moreover, we used lentiviral vectors to generate human cell lines that stably produce VLPs. Because VLPs can bind to the virus natural receptors, hence leading to entry into cells and viral antigen presentation, this platform could be used to develop novel vaccine candidates that are delivered intranasally.

**Highlights:**

- Identification of protein requirements for SARS-CoV-2 VLP production by transient transfection
- Lentiviral transduction to create cells stably producing SARS-CoV-2 VLPs
- Isolation of cell clones for the production of SARS-CoV-2 VLPs
- New putative platforms for vaccine development

## 1. Introduction

The severe acute respiratory syndrome coronavirus 2 (SARS-CoV-2), a recently emerged member of the *Coronaviridae* family, is responsible for the ongoing pandemic declared in March 2020 by the World Health Organization (WHO). Previous outbreaks of pathogenic coronaviruses occurred in 2002 and 2012 (SARS-CoV-1 and MERS-CoV, respectively). SARS-CoV-2 is relatively close to SARS-CoV-1 phylogenetically [1], yet is less harmful but more infectious with a higher spreading rate [2]. As a result of the rapidly spreading virus, novel vaccines were quickly developed [3]. Just a year after the pandemic emergency declaration by WHO, more than 100 COVID-19 vaccines candidates were in clinical stages of development, and close to 20 were approved [4]. Among the approaches undertaken are mRNAs and adenoviral vectors expressing the SARS-CoV-2 surface (S) protein [5]. These vaccines initially reported high effectiveness, *e*.*g*. 95% effectiveness for the BNT162b2 mRNA COVID-19 vaccine (Pfizer-BioNTech) [6], 94.1% for the mRNA-1273 vaccine (Moderna) [7], and 70.4% for the adenoviral vector ChAdOx1 vaccine (AZD1222; Oxford-AstraZeneca) [8]. However, the rapid evolution of the virus leading to the emergence of new variants of concern contributed to significant losses in vaccine effectiveness [9]. Regardless of virus evolution, injectable SARS-CoV-2 vaccines quickly lose effectiveness, consistent with the rapid reduction in vaccine-induced antibodies months after vaccination [10]. In particular, injectable vaccines elicit weak mucosal immunity in the airways. This is partly explained by low levels of local secretory IgA antibodies production, which is also linked to SARS-CoV-2 transmissions from vaccinated people [11]. Recently, progress has been made toward developing mucosal vaccines, delivered through intranasal and oral routes [12]. In addition to potentially improving effectiveness, mucosal delivery may facilitate large scale campaigns, due to the non-invasive route of administration.

As a relatively novel type of vaccine, virus-like particles (VLPs) represent an attractive solution to the challenges outlined above [13]. VLPs are formed through the natural self-assembly of viral proteins, hence mimicking native virions or subviral structures [14, 15]. Thus, VLPs present conformational epitopes, which are arranged repeatedly on the surface. Consequently, VLPs are particularly apt at inducing strong B cell responses in the absence of adjuvants [16]. In addition, the size of VLPs is favorable to uptake by antigen-presenting cells, leading to the induction of robust innate and adaptive immune responses, similar to the immunogenic properties of the natural virus [17-19]. The lack of genetic material makes VLPs non-infectious, and thus they can be manipulated in low-level biosafety laboratory settings [20]. Finally, VLP vaccines can rapidly cope with epidemic viral diseases because of the short time required for proceeding from design to expression [21]. Several expression systems have been described to generate VLPs, such as mammalian cell lines, bacteria, insect cell lines, yeast, and plant cells [22-24]. One important advantage of mammalian cells resides in the correct protein glycosylation pattern for VLPs produced in the mammalian cells [25]. Several teams have reported on the production of SARS-CoV-2 VLPs by transient transfection in mammalian cells or expression in insect cells, though there is no absolute consensus on which viral proteins are required to achieve efficient VLP assembly [26-30].

In this study, we report on the development of SARS-CoV-2 VLPs using transient transfection and stable transduction approaches. We demonstrate that the four structural proteins S, M, N and E are necessary and sufficient for efficient generation of VLPs. We use electron microscopy to further characterize those VLPs. Moreover, we successfully establish stable cell clones producing SARS-CoV-2 VLPs.

## 2. Materials and methods

### 2.1. Cell culture

HEK293T human embryonic kidney cells were cultured in Dulbecco’s modified essential medium (DMEM) (Fisher Scientific) supplemented with 10% fetal bovine serum (FBS) and penicillin/streptomycin (Hyclone). Cells were maintained in a 5% CO2 incubator at 37°C.

### 2.2. Plasmid construction and molecular cloning

The following SARS-CoV-2 structural protein-encoding plasmids were obtained from the Roth laboratory (University of Toronto): pDONR223_SARS-CoV-2-N, -S, -ORF3b, -ORF6, -ORF7a, -ORF8, as well as pDONR207_SARS-CoV-2-M, -E, -ORF3a, -ORF7a [31]. All SARS-CoV-2 protein coding sequences were cloned into the mammalian expression vector pEZY3, which was a gift from Yu-Zhu Zhang (Addgene plasmid #18672) [32] using the Gateway LR Clonase II Enzyme Mix kit (Invitrogen) as instructed by the manufacturer. 1 μl of each LR reaction was transformed into 50 μl of Library Efficiency DH5α competent cells (Invitrogen) using the heat-shock method. Transformed bacteria were plated on LB-agar dishes containing 50 μg/ml ampicillin and plates were incubated at 30°C overnight. Individual colonies were amplified and analyzed by restriction enzyme digestion and Sanger DNA sequencing of the inserts. To generate lentiviral vector constructs, SARS-CoV-2 S and M coding sequences were cloned into pLentiCMVPuroDEST (pLCPD), whereas E and N were cloned into pLentiCMVHygroDEST (pLCHD). pLentiCMVPuroDEST (w118-1) and pLentiCMVHygroDEST (w117-1) were gifts from Eric Campeau & Paul Kaufman (Addgene plasmids #17452 and #17454) [33]. SARS-CoV-2 coding sequences were inserted using the Gateway LR Clonase II Enzyme Mix kit (Invitrogen, CA), yielding plasmids pLCPD-S, pLCPD-M, pLCHD-E and pLCHD-N, respectively. Clonase reaction products were electroporated into competent *E. coli* (JM109 strain) and grown at 30°C. Plasmids were verified by restriction enzyme digestions and Sanger sequencing. Expression of SARS-CoV-2 S and N was verified by transfection into HEK293T cells followed by Western blotting of protein lysates (not shown). Expression of SARS-CoV-2 E and M proteins could not be verified due to the unavailability of commercial antibodies.

### 2.3. Generation of VLPs by transient transfection

HEK293T cells were seeded into 10-cm plates 24 h prior to transfection with pEZY3 plasmids expressing SARS-CoV-2 proteins using polyethyleneimine (PEI; PolyScience). For this, plasmid DNA (10 μg each) was diluted in 1 ml of serum-free, antibiotics-free cell culture medium. 45 μl of a 1 mg/ml aqueous PEI solution were added, followed by vortexing. Following a 10 min incubation on ice, the PEI-DNA solution was spread onto HEK293T cells plated the day before in a 10-cm tissue culture dish at approximately 50% confluence. Supernatants were replaced with complete medium the following day, and were collected 48 h post transfection by centrifugation at 3,000 rpm for 10 min at 4°C. Supernatants were filtered through 0.45 μm PVDF filters (Millipore), loaded on top of 20% sucrose-TNE cushions and ultracentrifuged for 90 min at 30,000 rpm at 4°C. The pelleted particles were recovered in TNE buffer [50 mM Tris-HCl, 100 mM NaCl, 0.5 mM EDTA (pH 7.4)]. Meanwhile, cells were washed with cold PBS, and then harvested and lysed with prechilled lysis buffer (10 mM Tris-HCl, pH 8.0, 1 mM EDTA, 1% Triton X-100, 0.1% Sodium Deoxycholate, 140 mM NaCl) supplemented with protease inhibitor cocktail (Pierce). Pelleted supernatants and cell lysates were immediately processed for Western blotting or frozen at -80°C.

### 2.4. Lentiviral vectors production and transductions

To produce lentiviral vectors encoding S, M, E, or N, HEK293T cells seeded in 10-cm plates at 80% confluency were PEI-transfected with 10 μg of pLCPD-S, pLCPD-M, pLCHD-E or pLCHD-N, 10 μg of the packaging construct pΔR8.9 and 5 μg of the VSV G-expressing construct pMD2G, as described previous [34]. The next day, supernatants were removed and replaced with fresh medium. Supernatants containing lentiviral vectors were harvested 2 days post-transfection and clarified by low-speed centrifugation.

HEK293T cells seeded in 6-well plates at 60% confluency were challenged with lentiviral vectors expressing combinations of S, M, E and/or N in the presence of 5 μg/ml polybrene (Millipore). 1 ml of each undiluted vector preparation was used for transduction of cells plated in one well, except for the simultaneous transduction with all 4 viral vectors, in which case we tried two conditions: 1 ml of each vector S/M/N/E (“SMNE1”), and 1 ml of each vector M/N/E mixed with 2 ml of the S vector (“SMNE2”). Supernatants were replaced with fresh medium the next day. One day later, cells were submitted to selection with 2 μg/ml puromycin (Gibco) and/or 100 μg/ml hygromycin (Enzo Life Sciences). Antibiotic selection was done for two days, which killed control untransduced cells.

To establish clonal cultures, transduced and selected SMNE2 cells were seeded in 96-well plates at 0.5 cells/well in 100 μl, using medium supplemented with 10% conditioned medium. The conditioned medium was obtained by collecting the supernatant of confluent HEK293T cells, then purifying it by centrifugation for 10 min at 3000 rpm followed by filtration through 0.45 μm PVDF filters (Millipore). Cell growth was monitored for one week. Wells that contained only one colony were selected and transferred to 10-cm plates. Purification of SARS-CoV-2 VLPs in supernatants was performed as before.

### 2.5. Western blotting

20 μl of VLPs or cell lysates were denatured using Laemmli sample buffer (0.125 M Tris pH 7.0, 4% SDS, 20% glycerol, 0.004% bromophenol blue,) and boiled at 95°C for 5 min. Proteins were separated by SDS-PAGE and then transferred to nitrocellulose membranes (Millipore). Membranes were probed with a 1:1000 dilution of anti-SARS-CoV-2 S mouse monoclonal primary antibody (1A9, GenTex) or a 1:1000 dilution of anti-N rabbit polyclonal primary antibody (GenTex). HRP-conjugated secondary anti-mouse and anti-rabbit IgG (Cell Signaling Technology) were diluted at 1:10,000. Membranes were scanned using the multiplex scanning method on the Azure Biosystems scanner (software version 1.2.1228.0) according to the manufacturer’s manual.

### 2.6. Electron microscopy

Supernatants were purified using low-speed centrifugation for 10 min at 3,000 rpm, followed by filtration through 0.45 μm PVDF filters (Millipore). Filtrates were fixed with 1:10 volume of 37% formaldehyde for 10 min at room temperature, and the samples were then loaded on top of 20% sucrose-TNE cushions and ultracentrifuged for 3 h at 25,000 rpm at 4°C in a Sorval WX Ultra 100 ultracentrifuge. The final pelleted particles were recovered in 100 μl of 0.2 M sodium cacodylate buffer, pH7.4 (Electronic Microscopy Science). 10-20 μl of this sedimented material were incubated on carbon and formvar-coated grids for 30 sec. The grids were negatively stained with 4% Uranyl Acetate for 30 sec then air dried for a few minutes before observation with a transmission electron microscope (Philips EM208S Transmission Electron Microscope at 80 kV).

## 3. Results

### 3.1. Generation of SARS-CoV-2 VLPs in human cells using transient transfection

To define the minimal SARS-CoV-2 structural proteins required to generate SARS-CoV-2 VLPs, we first transfected HEK293T cells with plasmid vectors expressing each of the four structural proteins, *i*.*e*. S, M, E, or N, either alone or in combination. VLPs were purified using low-speed centrifugation followed by filtration. The particles were concentrated using 20% sucrose cushions followed by immunoblotting (Fig. 1A). We detected the expression of S and N proteins in whole cell lysates (WCLs) and the ultracentrifuged fraction (UCF). At the time of this work, no antibodies were available for M and E. As shown in Fig. 1B, S and N were detected in WCLs in the presence or absence of other structural proteins. The anti-S antibody detected two bands at around 200 kDa in cells, similar to what was observed by other authors [35], as well as the S2 furin cleavage product [36] as expected for this antibody. High amounts of both S and N were detected in the UCF pellets upon co-transfection of SARS-CoV-2 N, S, E and M. In contrast, S was found in smaller amounts in UCF fractions of cells transfected with S alone or in combination with one or two other structural proteins (Fig. 1B). Similarly, N was found in smaller amounts in the UCF pellets of cells transfected with N+S, N+S+E or N+S+M, and was not seen at all when transfected alone. Altogether, this result suggests that all four structural proteins S, M, N and E are required for the efficient assembly and egress of SARS-CoV-2 VLPs. However, both S and N seemed to be present in particulate form in the supernatants of cells in conditions in which the formation of true SARS-CoV-2 VLPs is not expected to occur, for instance upon transfection of S alone. This underlines the challenges of segregating true VLP assemblies from protein aggregates or other forms of particulate materials.

**Fig. 1.**
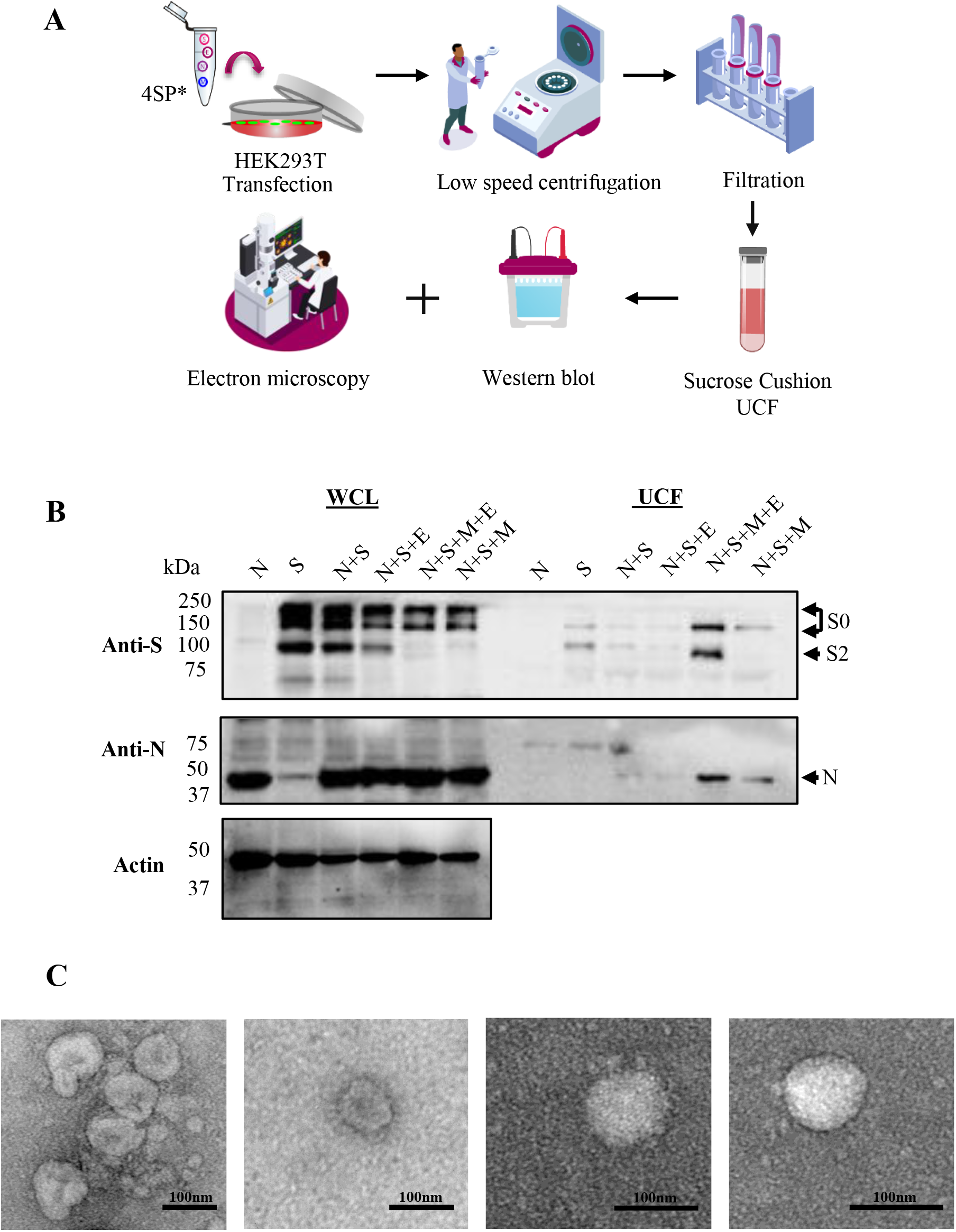
Production of SARS-CoV-2 VLPs by transient transfection. (A) Schematic outline of SARS-CoV-2 VLPs generation in HEK293T cells transfected with SARS-CoV-2 structural protein-expressing plasmids. (B) Western blot analysis of VLPs in supernatants of transfected cells. SARS-CoV-2 proteins present in whole cell lysates (WCLs) and ultracentrifuged supernatants (UCF) 48 h after transfection with the indicated plasmids, and detected using anti-S and anti-N antibodies. Protein size markers are shown. This experiment is representative of two independent replicates. (C) Electron microscopy detection of SARS-CoV-2 VLPs. Supernatants of transfected HEK293T cells were collected 48 h after transfections, purified, fixed, concentrated by ultracentrifugation and processed for EM analysis. Scale bar = 100 nm.

We used transmission electron microscopy (EM) to confirm the presence of virion-like VLPs in the supernatant of cells transfected with the four structural proteins. Several protocols were tested for the fixation, concentration and staining of samples putatively containing VLPs. We settled on a protocol in which supernatants are clarified by low-speed centrifugation and filtration, then fixed with formaldehyde prior to concentration by ultracentrifugation, and UCF pellets are deposited on grids and negatively stained. As shown in Fig. 1C, particles with a SARS-CoV-2 virion-like morphology were found that displayed typical corona-like spikes. These particles were about 100 nm in diameter, spikes included (Fig. 1C), which is consistent with SARS-CoV-2 particles. These observations strongly suggest that SARS-CoV-2 VLPs were generated through this transient transfection approach.

### 3.2. Role of SARS-CoV-2 accessory proteins in VLP production

The SARS-CoV-2 ∼30 kb genome encodes accessory proteins, the exact number of which is still under debate [37]. These accessory proteins have important roles in virus infectivity and pathogenicity [38]. For instance, ORF3a and ORF7 promote immune escapes by downregulating MHC expression at the surface of infected cells [39]. It is conceivable that some accessory proteins would indirectly increase SARS-CoV-2 VLP production. For instance, ORF8 is known to increase viral replication through the modulation of endoplasmic reticulum functions [40]. To investigate the possibility that SARS-CoV-2 accessory proteins increase VLP production, we transfected HEK293T cells with the four structural proteins alone or in combination with accessory proteins. As shown in Fig. 2, none of the accessory proteins tested increased S and N expression in transfected cells. In fact, ORF3a/b, ORF6 and ORF7a/b seemed to decrease N protein expression. Analysis of the ultracentrifuged supernatants showed that the cells transfected with the four structural proteins and no accessory proteins produced higher levels of VLPs than in the other conditions tested. These data suggest that the four SARS-CoV-2 structural proteins are sufficient for the efficient formation of VLPs.

**Fig. 2.**
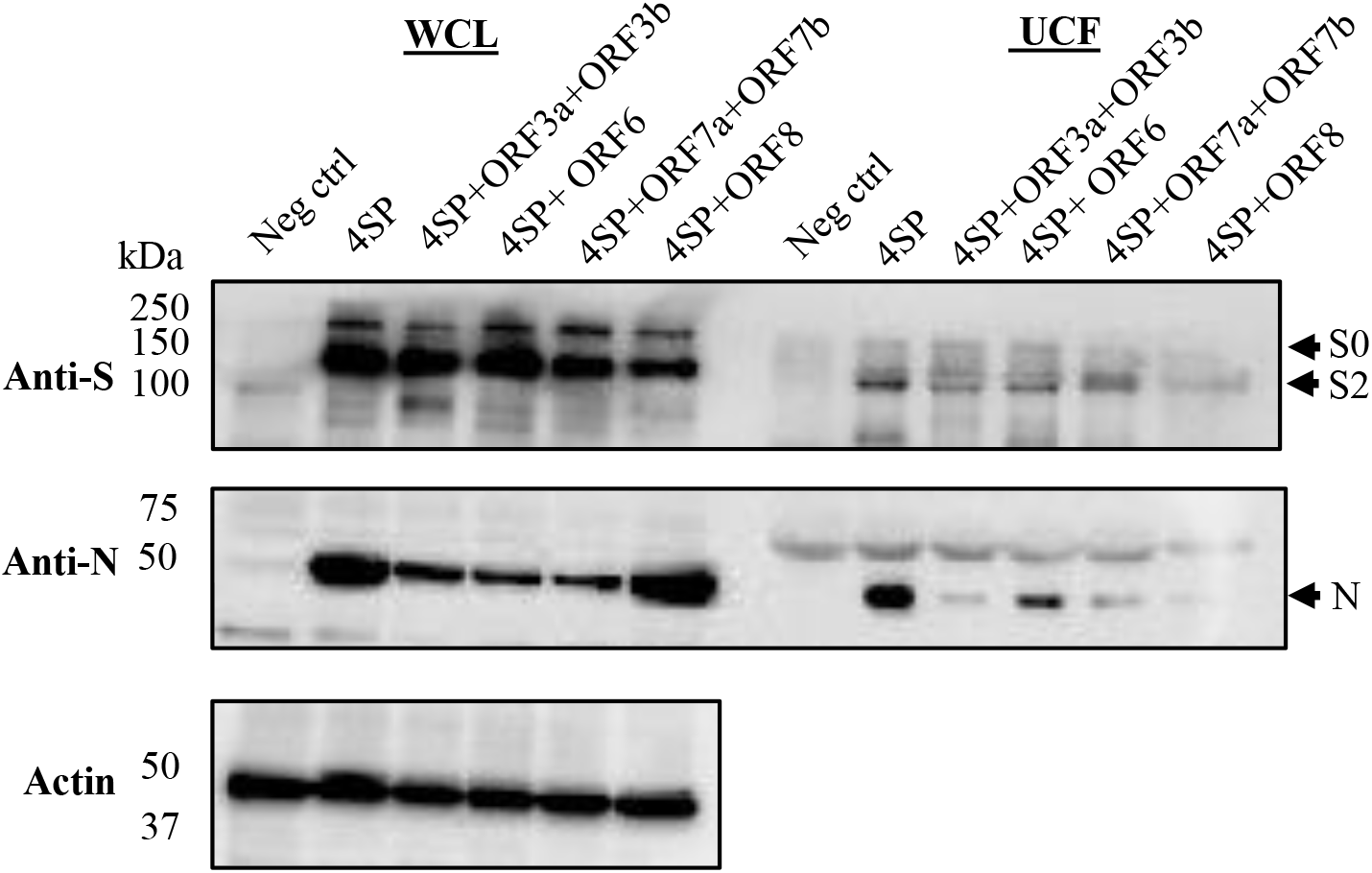
Characterization of minimal protein requirements for efficient SARS-CoV-2 VLPs production. Western blot analysis of SARS-CoV-2 S and N proteins in WCLs and in ultracentrifuged supernatants (UCF) 48 h after transfection of the plasmids expressing the indicated proteins. 4SP (“4 structural proteins”) stands for a mixture of S, M, E and N-expressing plasmids.

### 3.3. Efficient production of SARS-CoV-2 VLPs using lentiviral vectors

As described above, we were able to generate SARS-CoV-2 VLPs by transient transfection. To facilitate VLP production, we explored the possibility of creating HEK293T cell lines stably expressing VLPs through lentiviral transduction. This approach would allow cells to continuously release VLPs into the cell culture supernatant, eliminating the need for transient transfection of plasmids. As summarized in Fig. 3A, we first generated S, M, E and N lentiviral vectors by transfecting cells with appropriate plasmid combinations. The vectors encoding SARS-CoV-2 S and M also carried puromycin resistance, whereas the vectors encoding N and E carried hygromycin resistance. HEK293T cells were transduced with the four lentiviral vectors simultaneously, or with single vectors. We were concerned that the transduction of S would be less efficient owing to the large size of its coding sequence. Thus, we tried two different transduction conditions for the quadruple transduction: either an equivalent volume of all four lentiviral vectors (SMNE1) or twice as much of the S vector, relative to the four other vectors (SMNE2). Whole cell lysates were prepared, and supernatants were purified and concentrated as previously, yielding UCF fractions that contained particulate materials that sedimented through the sucrose cushion. Both WCLs and UCF samples were then analyzed by immunoblotting. As shown in Fig. 3B, S and N were detected in the WCLs of cells transduced with the four structural proteins or with the relevant individual proteins, as expected. Of note, we did not see any difference in S expression between SMNE1 and SMNE2. Surprisingly, both S and N were readily detected in UCF pellets of cells transduced with a single lentiviral vector, in addition to being present in the supernatants of cells transduced with the four structural proteins. Presumably, the particulate materials purified from cells transduced with N or S alone are either aggregates and/or exosomes containing the proteins. But because the signals in cells transduced with one or 4 vectors were of similar intensities, this Western blotting analysis did not allow us to demonstrate the presence of “true” VLPs.

**Fig. 3.**
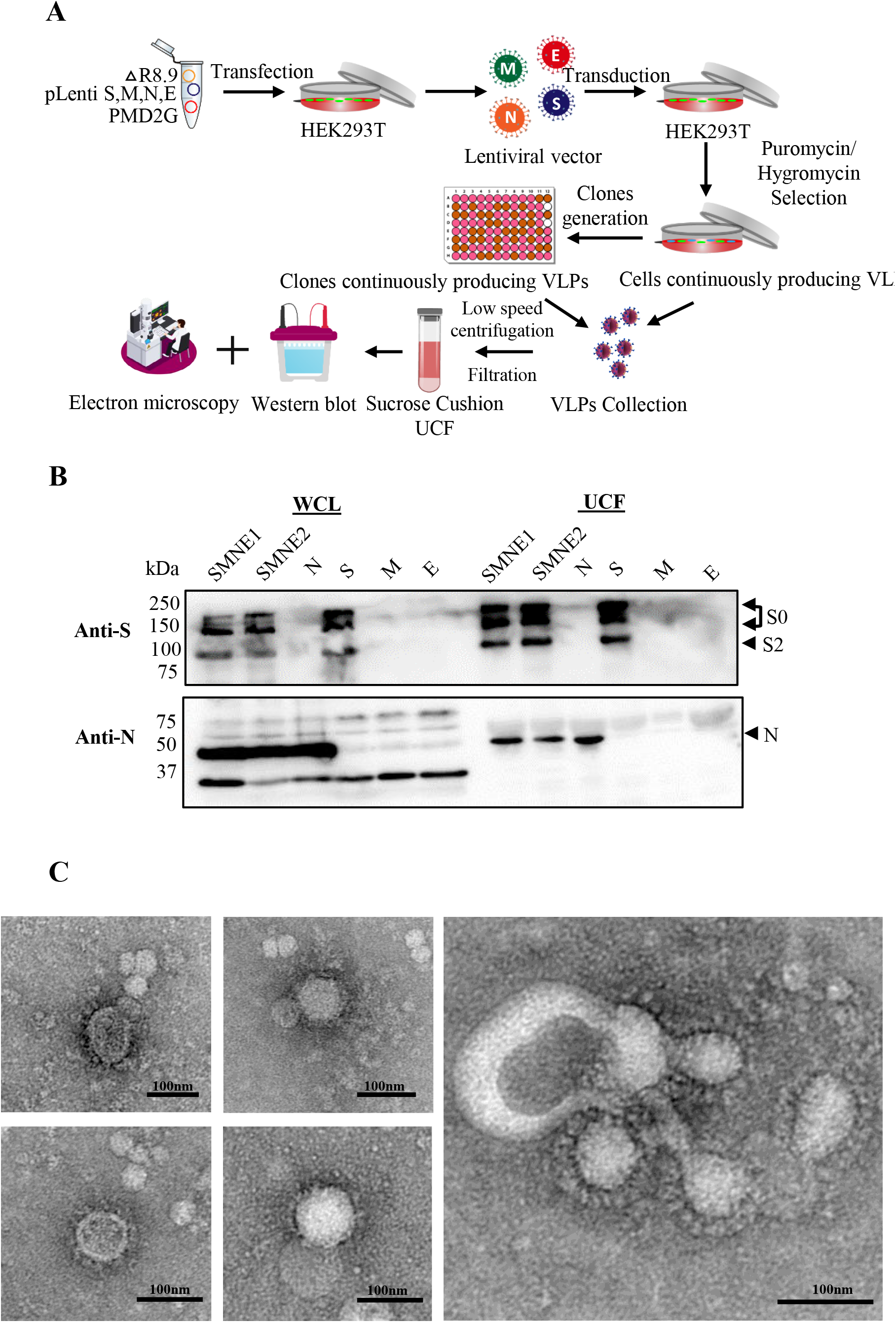
Generation of SARS-CoV-2 VLPs using lentiviral transduction. (A) Schematic outline for the generation of cell populations and cell clones producing SARS-CoV-2 VLPs. (B) Western blot analysis of SARS-CoV-2 proteins S and N in WCLs and ultracentrifugated supernatants (UCF) from cells transduced with lentiviral vectors expressing the indicated SARS-CoV-2 proteins. SMNE1 and SMNE2 refer to differences in the amount of the S vector relative to the other vectors (see 2.4). Protein size markers are shown. This experiment is representative of two independent replicates. (C) Electron microscopy detection of SARS-CoV-2 VLPs. Shown are EM images of putative SARS-CoV-2 VLPs produced by HEK293T cells transduced with S, M, N and E. The two pictures at the top-left, and the one at the bottom-left, show particles from “SMNE1” cells, whereas the two other images are from “SMNE2” cells. Scale bar = 100 nm.

To confirm the formation of VLPs in cells transduced with the four SARS-CoV-2 structural proteins, we analyzed the UCF fractions using EM. As shown in Fig. 3C, we found particles that resembled SARS-CoV-2 virions in shape and size with the presence of distinctive spike protein. We did not find similar structures in cells transduced with single vectors, though the observations were too infrequent for us to perform a quantitative analysis. Therefore, these data suggest that we successfully generated mammalian cell lines stably producing SARS-CoV-2 VLPs.

### 3.4. Generation of cell clones stably producing SARS-CoV-2 VLPs

For industry-scale applications of transduced mammalian cell lines continuously producing SARS-CoV-2 VLPs, it is desirable to isolate cell clones that produce high amounts of VLPs. For this, single cell cultures of HEK293T cells transduced with the four structural proteins were initiated by limiting dilution of the cells transduced with all four lentiviral vectors (“SMNE2”). Surviving cell populations were amplified, and we analyzed the presence of N and S in purified, ultracentrifuged supernatants by immunoblotting followed by EM analysis. As shown in Fig. 4A, all the analyzed clones expressed S and N, but significant variations were observed, especially for S. EM analyses (Fig. 4B) showed the presence of VLPs similar in shape and size to those found before. Thus, these data suggest that most or all cells transduced with the four structural proteins produce SARS-CoV-2 VLPs, but that analysis of individual cell clones is useful to identify the most promising ones for downstream applications.

**Fig. 4.**
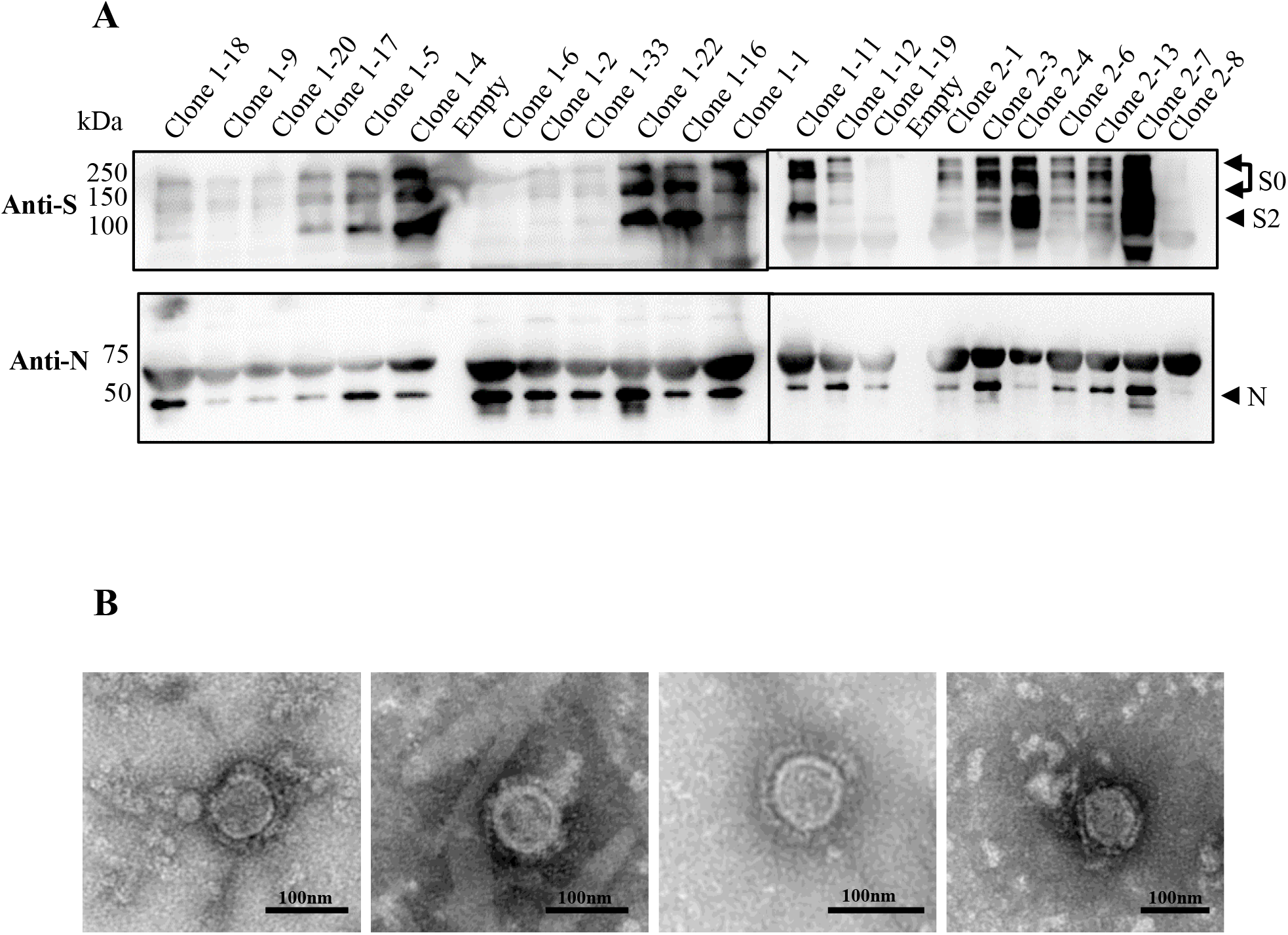
Generation of cell clones stably producing SARS-CoV-2 VLPs. (A) Western blot analysis of SARS-CoV-2 proteins S and N present in ultracentrifugated supernatants of 22 cell clones derived from HEK293T cells transduced with S, M, N and E (“SMNE2” cells) as shown in Fig. 3. Protein size markers are shown. (B) Morphological analysis of SARS-CoV-2 VLPs. Shown are EM images of putative SARS-CoV-2 VLPs found in the supernatants of HEK293T clones (from left to right: 1-22, 1-16, 2-13, 2-7). Scale bar = 100 nm.

## 4. Discussion

Virus-like particles (VLPs) are nanoscale structures made up of self-assembled viral proteins, mimicking the native virus but lacking the genetic material so they are not capable of productively infecting the host cell [41]. VLPs have gained increasing interest as immunogens capable of eliciting both mucosal and cellular immune responses [42]. Their repetitive antigenic structure enables them to induce strong T-helper response [43]. VLPs are also prone to activate dendritic cells (DCs) which are essential players in the initiation of an immune response through the capture and processing of antigens, delivering them to secondary lymphoid organs and providing co-stimulatory signals [44]. VLPs produced in mammalian cells offer additional benefits, including the possibility of VLP surface proteins glycosylation [45]. In the present study, we generated SARS-CoV-2 VLPs by co-expressing four SARS-CoV-2 structural proteins in HEK293T, a mammalian cell line commonly used to produce biological reagents such as monoclonal antibodies. We first transfected HEK293T cells with vectors expressing S, M, E, or N, respectively, as well as combinations of these proteins. Other studies had done similar work, and they detected the presence of viral proteins in supernatants even when just 1, 2 or 3 proteins were transfected [26]. In contrast, our data support the notion that all four structural proteins are essential for the efficient release of VLPs. It is likely that the discrepancy could be explained by a better purification procedure in our study.

Our results suggest that we created HEK293T cell populations stably expressing VLPs through lentiviral transduction. This approach may allow cells to continuously release VLPs into the cell culture supernatant, facilitating large-scale VLP production. Future experiments should include a more detailed characterization of VLPs produced in this way, such as time-dependent analysis of VLP production over extended periods of time. We further isolated cell clones that seemingly produce high amounts of VLPs. Such clonal populations might prove useful for downstream industrial applications. However, in the context of these VLPs produced through lentiviral transduction, Western blotting was insufficient to prove true VLP assembly, since we were able to sediment SARS-CoV-2 proteins from the supernatants of cells transduced with S or N alone. Why was this problem encountered in the context of transduced but not transfected cells, is presently unclear. It will be important to develop methods that allow for more efficient VLP purification. However, using electron microscopy we observed the presence of viral particles displaying typical S protein projections that are a hallmark of coronaviruses, in the supernatant of both transfected and transduced cells, as well as clonal populations of transduced cells. Those particles were about 100 nm diameter, which is comparable to wild-type virions.

Previous studies had shown that some accessory proteins such as ORF3b, ORF6, ORF7a and ORF8 are important IFN-I antagonists, leading to an impairment in the host immune response [37]. It is conceivable that some of these proteins would also be important for VLP egress, perhaps by inhibiting innate immune responses targeting these stages of viral replication. To test this hypothesis, we co-transfected HEK293T cells with the four structural proteins in combination with selected accessory proteins. Our data showed that no additional proteins other than the four structural proteins are required for SARS-CoV-2 VLPs assembly and release.

The term “VLPs” is the subject of confusion, as it is used to describe not only assemblies of proteins from the virus of interest, as in this study, but also the incorporation of a single viral protein (typically a surface protein) in lipid droplets or heterologous viral particles [46], and even assemblies of a single type of viral protein that may not have a true viral structure at all. For instance, the company Medicago has developed a plant-based vaccine for SARS-CoV-2 by expressing the S protein in *Nicotiana benthamiana* plants; though the resulting immunogens are called “VLPs” in the literature, they may simply be S aggregates [47]. In the context of SARS-CoV-2 VLPs used as vaccines, the presence of multiple SARS-CoV-2 viral proteins (*i*.*e*. multiple antigens) as in the current study might result in increased effectiveness, compared to current vaccines that typically include only one, and might also be less susceptible to loss of effectiveness due to virus evolution. It should also be pointed out that the VLPs containing all four SARS-CoV-2 structural proteins are expected to fuse into target cells in a manner similar to actual viruses. This feature might translate into increased immunogenicity compared to antigens that must be actively internalized by antigen-presenting cells. From the perspective of fundamental virology research, the systems described here could constitute useful tools for the study of not only assembly and egress but also virus entry.

## 5. Conclusions

In this study, we successfully generated SARS-CoV-2 VLPs from mammalian cells, either transiently or through stable transduction. These VLPs presumably contained all four structural proteins and thus closely resemble SARS-CoV-2 virions, which may translate into higher mucosal immunogenicity compared with vaccines composed of only one viral protein. However, VLPs immunogenicity remains to be tested, and VLP purification needs to be improved. In addition to the development of nasally-administered vaccines for mucosal immunity against coronaviruses, the tools presented here could also be useful to the study of virus assembly, egress and entry.

## Acknowledgments

We are grateful to Dr Frederick Roth (University of Toronto) for sharing SARS-CoV-2 plasmids.

